# The end of protein structure prediction: Improving prediction accuracy in chimeric proteins by windowed multiple sequence alignment

**DOI:** 10.1101/2024.10.06.616858

**Authors:** Sanketh Vedula, Alex Bronstein, Ailie Marx

## Abstract

AlphaFold2 has predicted the structures of almost every known protein. A simple means to create proteins beyond those found in nature, is by unnaturally fusing together two known proteins. Here we demonstrate that dependence on multiple sequence alignment, limits the success with which AlphaFold and ESMfold capture such chimeric forms of otherwise well predicted, individual, proteins. Specifically we show that peptides are predicted with significantly reduced accuracy when added to the terminal *ends* of scaffold proteins. Appending the multiple sequence alignment for the individual peptide tags to that of the scaffold protein often restores prediction accuracy.

## 1 Introduction

Proteins have evolved over billions of years, accumulating sequence variations over time. Recognizing that coevolution of positions distant in the protein sequence is indicative of proximity in the protein structure revolutionized attempts to predict structure from sequence [5]. This concept has remained a cornerstone of state of the art, deep learning, prediction methods where the same key information is extracted as subtle signals from alignments of large numbers of sequences – the so called multiple sequence alignment (MSA) step [7]. Clustering or shallowing the MSA has facilitated greater conformational sampling by AlphaFold2 [2, 17]

Here we demonstrate that using default MSA parameters is likely to give low accuracy prediction of chimeric proteins and show that the prediction accuracy of the individual protein parts is restored when combining the MSA information for each region. This problem has immediate practical importance as evidenced by the recent use of AlphaFold2 in chimeric immunogen design [15]. Our results suggest that AlphaFold [7] and ESMFold [6] have very limited inductive capabilities.

## 2 Experimental Setup

### Structure prediction

We compare three protein structure prediction algorithms, AlphaFold2 [7], AlphaFold3 [1], and ESMFold3 [6]. AlphaFold2 uses co-evolutionary information extracted from MSA and produces a point-wise, and pair-wise representations (effectively “a contact map”) using the `Evoformer` module; these representations are then decoded into 3D coordinates via the `Structure` module. AlphaFold3 also uses MSAs to construct pair-wise features, which are used to condition a diffusion model to produce 3D coordinates of the folded structure. ESM family of approaches [12, 11, 6], on the other hand, do not perform explicit MSAs. Instead, they first training a protein language model to solve a masked language modeling task in the style of BERT [3]. A separate model is trained to predict the contact maps from the learned representations of the ESM language model. We use ColabFold [10] to perform AlphaFold2 predictions. For the ESMFold3, we use the recently-released ESM3 language model [6], and its structure prediction head. For ESMFold3, we considered both iterative and `argmax` decoding as recommended in [6]; we set the iterative decoding version as the main baseline because of its better accuracy. We obtained AlphaFold3 predictions from https://alphafoldserver.com^1^.

### Dataset creation

We create a large set of chimeric proteins by adding the sequences of short peptides at the `N` and `C` terminal of scaffold proteins, `SUMO2`, `GST`, `GFP` and `MBP`. These proteins are commonly employed as inert fusions to aid expression, purification and cellular tracking of target proteins. We note that the scaffold sequences used correspond to that found in the crystal structures, `2IYD:B`, `1PKW`, `2B3P` and `1MPB` respectively, meaning that the `SUMO` and `MBP` proteins were truncated at the `N` terminus with respect to the native protein. The peptide tags were selected from a recent benchmark testing the performance of AlphaFold2 on peptide structure prediction [9]. We first perform a redundancy filtering on the tag sequences, selected peptides with a low overall RMSD (<1Å) between the experimentally determined structure. This results in 34 tags, which are listed in Table A1. Chimeric proteins were created by the addition of peptide tag sequences to the `C` and `N` terminus, individually, of the scaffold proteins. A small and flexible `GLY-SER` linker was inserted between the protein parts to alleviate any potential steric constraints in the concatenated sequences.

### Chimeric protein structure predictions

We run predictions using AlphaFold2 and ESMFold3 for all combinations of aforementioned 4 scaffolds with 34 tags (see Tab. A1), attached once at `N` and `C` terminals, resulting in total 252 unique sequences. We assess the accuracy of the tag prediction was assessed by calculation of the RMSD between the experimentally determined peptide structure and the peptide sequence region of the chimeric protein. Because Alphafold3 is a limited-access server, we run Alphafold3 predictions only for the examples presented within the Fig. 1.

**Figure 1:**
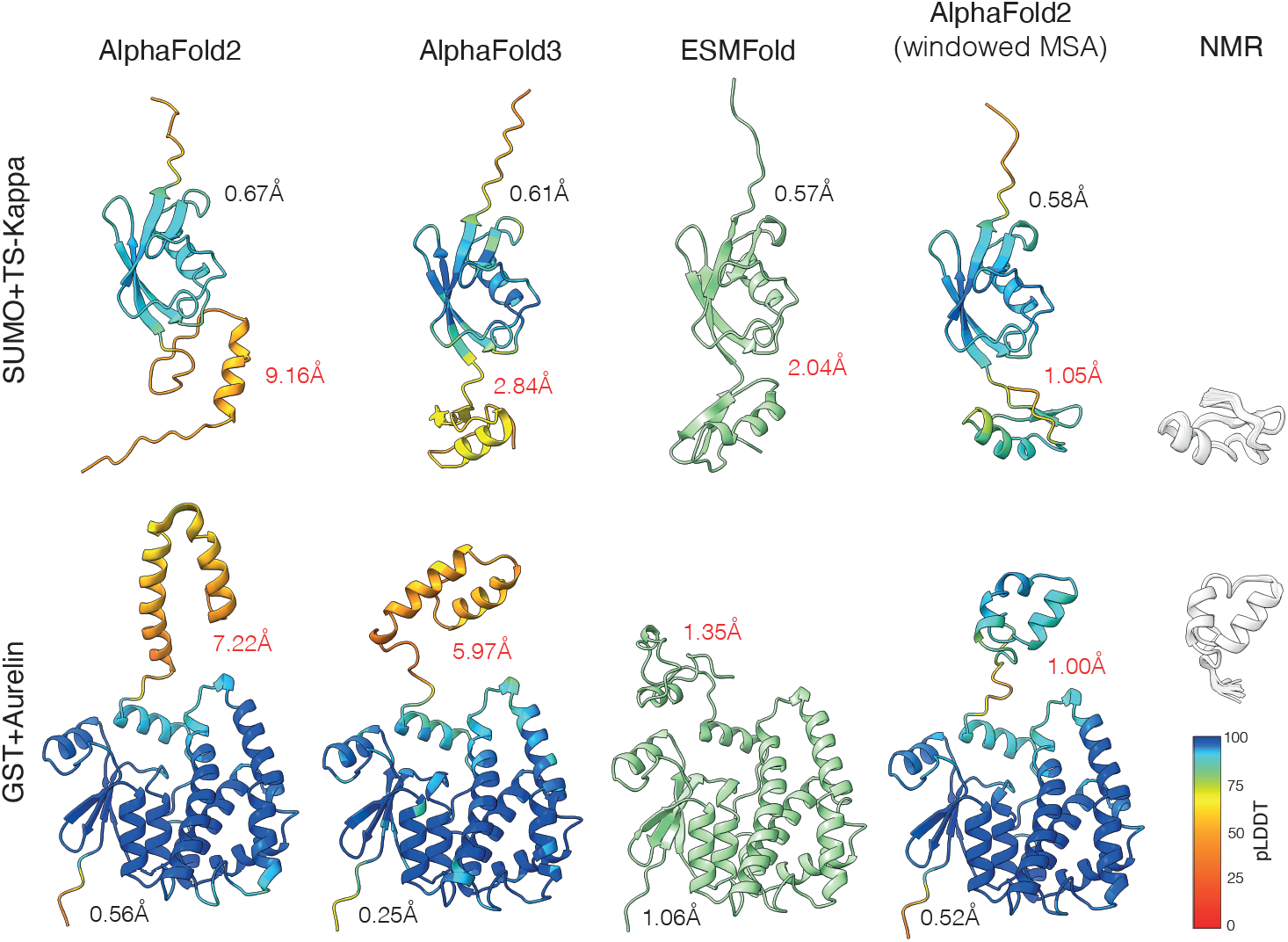
Protein structure prediction algorithms fail to predict the structure of the tag accurately in the scaffold context. AlphaFold2 with the proposed windowed MSA predicts accurately. Shown left-to-right are predicted structures of two synthetic chimeric sequences (first row: `1TSK+SUMO`; second row: `GST+2LG4`) by AlphaFold2, AlphaFold3, ESMFold, and AlphaFold2 with the proposed windowed MSA technique. AlphaFold-predicted structures are colored by the pLDDT confidence. The rightmost column shows NMR structures of the attached peptide tags. Prediction accuracy is reported in terms of RMSD separately for the scaffold (in black) and the tag (in red). While the scaffolds are invariably predicted accurately, tag structures are mispredicted by the MSA-based AlphaFold and predicted very accurately with the proposed “fix”. ESMFold shows a somewhat better prediction of the tag, but it is still significantly worse than the accuracy of the tag predicted without the scaffold context.

## 3 Results

### Peptide structure prediction

Fig. A4 presents a comparison of AlphaFold2, ESMFold-iterative, and ESMFold-argmax predictions on the peptide structure prediction benchmark introduced in [9]. We observe that AlphaFold2 makes more accurate predictions compared to ESMFold. Out of the 394 tags that were benchmarked, AlphaFold2 attains an accuracy of <1Å RMSD on 34 tags (Table A1), whereas ESMFold-argmax and ESMFold-iterative attain <1Å RMSD on 18 and 21 tags, respectively.

### AlphaFold2 and ESMFold fail to accurately predict tags in context of a scaffold

We filter the tags that were predicted accurately (RMSD <1Å) by AlphaFold2 and ESMFold, respectively, and measure the accuracy of these “good tags” when predicted in context of a scaffold at both terminals. Representative results are presented in Fig. 1, and they demonstrate that both AlphaFold2 and ESMFold predict tags poorly when predicted in the context of a scaffold protein. Fig. A1 presents a tag-level breakdown of the ratio of the RMSD of the tag when predicted in context with respect to the RMSD when predicted in isolation. We can observe that for most tags, this ratio is greater than 1, indicating that adding a context worsens the prediction of the tag’s structure.

### This problem persists in AlphaFold3

Due to the limited accessibility of AlphaFold3, we were unable to perform an extensive quantitative analysis. However, we obtained AlphaFold3 predictions for a handful of structures, some of which are represented in Fig. 1. While AlphaFold3 generally exhibits superior prediction accuracy, it seems to fail to accurately predict tag structures in a bigger context.

**Windowed MSA.** We hypothesize that the inability of AlphaFold to accurately predict a tag when presented within a scaffold context – despite accurately predicting the tag in isolation – is due to their reliance on MSA. When the tag region is combined with the scaffold, forming a chimeric sequence, the MSA does not adequately represent the tag region because of the chimeric nature of the sequence. As a result, the tag is not preferentially treated during alignment, leading to inaccurate modeling when the tag is within the context.To address this issue, we propose augmenting the MSA of the chimeric protein sequence by incorporating the MSA of the tag region alone. By integrating the standalone MSA of the tag into the chimeric sequence’s MSA, we aim to ensure the tag region is properly represented during alignment. We then provide this augmented MSA to AlphaFold2 as input and perform the predictions [10, 7].

### AlphaFold2 with windowed MSA predicts the tags accurately in scaffold context

We run the AlphaFold2 with windowed MSA predictions for all scaffold-tag combinations by running AlphaFold2 directly on the updated MSA using ColabFold [10], and compare it to the predictions obtained with AlphaFold2 running on the standard MSA. Fig. 3 presents the results of the ratio of the RMSD of obtained by AlphaFold2 using the standard MSA with respect to the RMSD obtained using the windowed MSA technique, both measured at the tag. We observe that this ratio is > 1 in most cases (252 out of 272 predictions) implying that applying the windowed MSA “fix” allows predicting the terminal tag regions accurately. A visual depiction of the predictions obtained from AlphaFold2 and AlphaFold2 with windowed MSA is presented in Fig. 2, along with the regional MSA coverage. We note that not only does windowed MSA improve the accuracy of the prediction, but it also improves the pLDDT confidence score output by AlphaFold2.

**Figure 2:**
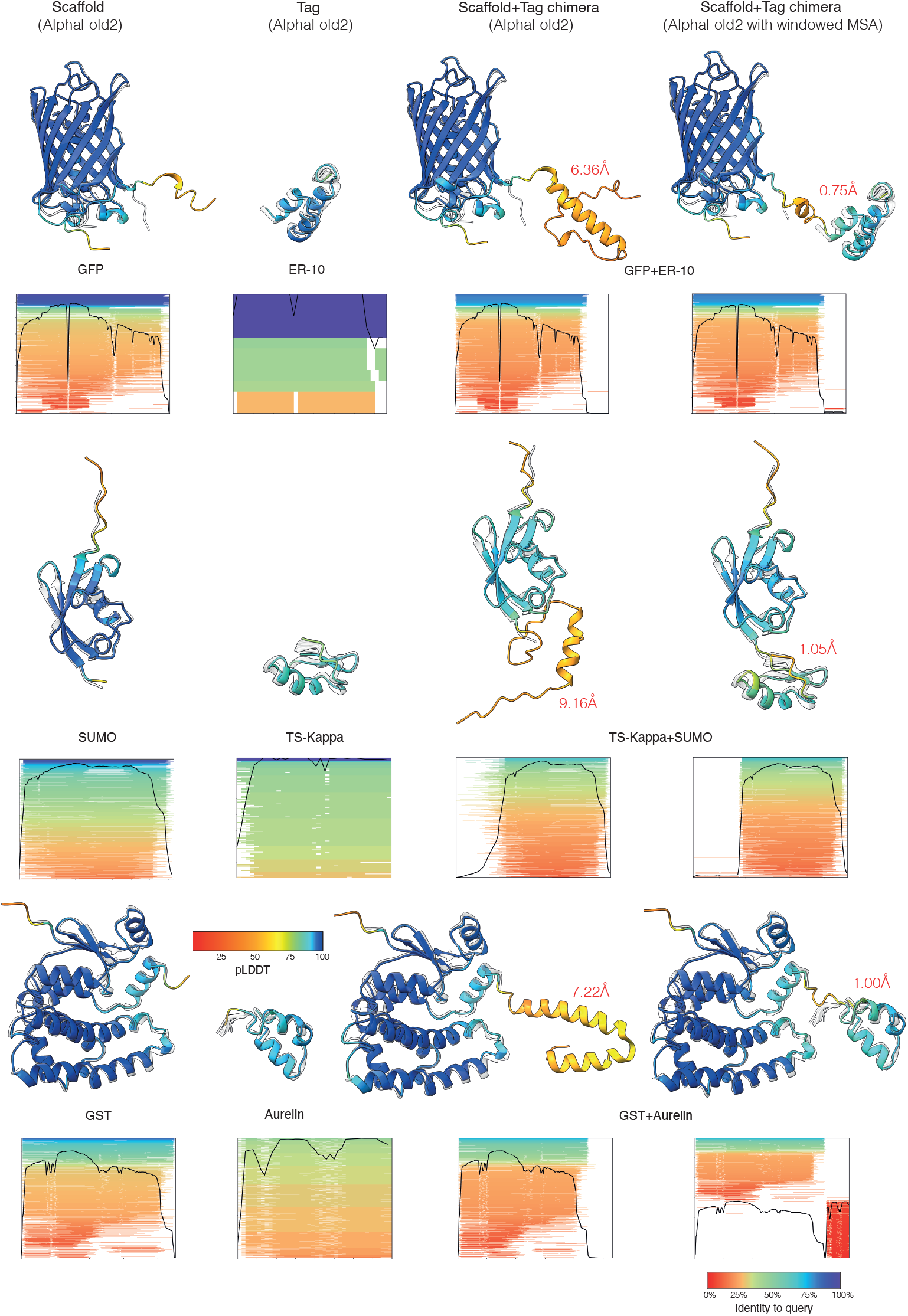
AlphaFold2 predicts exquisitely accurate structures for both the scaffolds and the tags individually wherever MSA coverage is high, and fails to predict the tag region in the chimeric sequence due to poverty of the MSA coverage. The proposed fix increases the coverage in the tag region which results in accurate structure prediction. Depicted left-to-right are the predicted structures for individually a scaffold and a tag, their combination into a chimeric sequence, and an improved prediction of the chimera using the proposed windowed MSA “fix”. Predicted structures are colored by the pLDDT confidence and superimposed on the experimentally determined structures (transparent white). RMSD of the tag prediction is reported in red. Below each predicted structure the MSA coverage is shown sorted and colored by identity to the query sequence. `GFP+1ERP`, `1TSK+SUMO` and `GST+2LG4` chimeras are shown.

**Figure 3:**
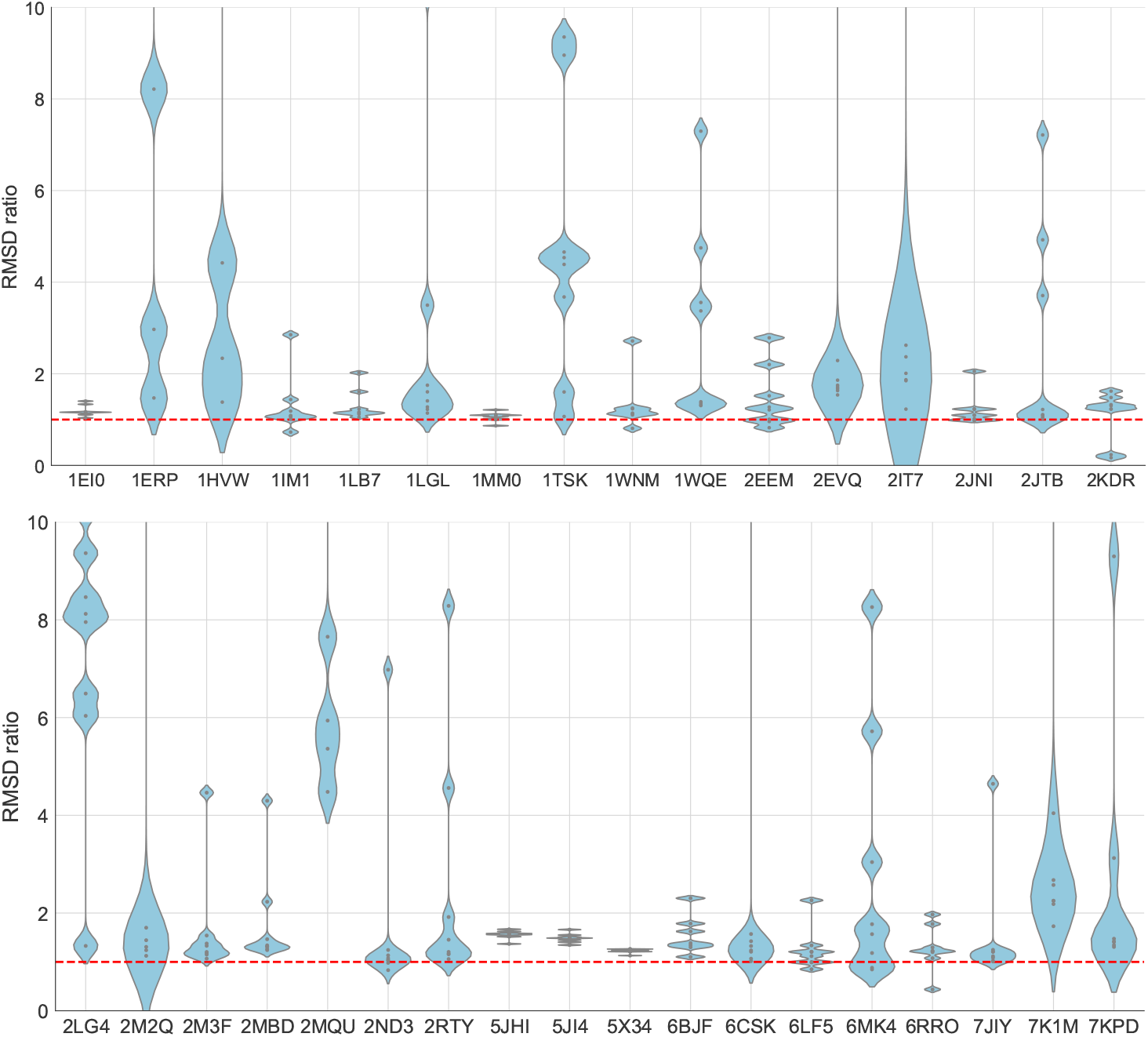
Windowed MSA + AlphaFold2 (AF2) accurately estimates the tags compared to AF2 in the context of a scaffold. Depicted are the ratios of the RMSD of AF2 with respect to the RMSD of Windowed MSA+AF2 per tag. We see that in most cases the ratio is greater than 1 (above the red dashed line) indicating that Windowed MSA + AF2 results in lower error than AF2.

## 4 Conclusion

Deep learning techniques combined with the exponential growth of available compute power due to the development of general-purpose GPUs caused tectonic shifts in various domains of science and engineering. The earliest moment of such a shift can be traced back to AlexNet beating all competing visual recognition algorithms on the ImageNet with a large margin [8]. Unlike its competitors invariably based on “hand-crafted” features constructed based on human understanding of the data (natural images in the case of ImageNet), AlexNet learned the representation entirely from the raw input data – a hallmark capability of deep learning^2^.

DeepMind’s AlphaFold 1 [13] halving the prediction error on the 2018 CASP benchmark [14] has been frequently equaled to the “AlexNet moment” in structural biology. However, besides the huge impact of both breakthroughs on their respective fields, this is a largely false equivalence. AlexNet and, to a larger extent, later neural network architectures such as visual transformers [4] are, in essence, glorified interpolators trained in a straightforward fully-supervised regime on massive amounts of labeled data and achieving its remarkable inductive capabilities – with very few hand-crafted inductive biases – due to the relative smoothness and low dimensionality of the image manifold.

In sharp contrast, the general protein sequence-to-structure relation is very likely much less regular in comparison to images, and the amounts of structural data available in the entire PDB are insufficient by many orders of magnitude to make the sequence-to-structure supervised learning feasible. Instead, in the heart of AlphaFold lies the fundamental discovery of the co-evolution of contacting residues [5]. Consequently, given enough evolutionary-related sequences, the analysis of correlated mutations by means of standard multiple sequence alignment (MSA) techniques allows to estimate residue contact maps. The task of reconstructing the protein structure given such maps as an input is a significantly easier task that can be addressed with deep learning techniques trained on the relatively modest (about 150K structures) training set available from the PDB. Without derogating from its revolutionary impact, AlphaFold does not learn the laws of protein folding; it rather “only” learns how to convert contact maps into atom coordinates based on a powerful hand-crafted inductive bias.

The importance of wide contexts (contacting residues are typically non-local in the sequence) for protein structure prediction has been demonstrated by protein language models. Sequence representations such as ESM [12, 11, 6] trained, similarly to natural large language models, with the task of sequence inpainting on the UniProt sequence data, have been shown to lead to accurate structure prediction with a relatively small and simple model trained on top of the sequence embedding [11, 6]. Similarly to AlphaFold, ESMFold neither learns how to fold proteins – it essentially “only” learns how to perform MSA using high-quality sequence embeddings, and in doing so it relies on the tens of millions of sequence records (lacking experimental structure data!) made available thanks to the ground-breaking commoditization of DNA sequencing techniques in the 90’s[16].

The results presented in this paper bring us very close to claiming that AlphaFold is as good as the multiple sequence alignment on which it relies. Practically, this implies that while it will achieve remarkable accuracy in predicting structures of evolution-produced sequences with rich phylogenetic information (high MSA coverage), the prediction of completely de novo sequences is hopeless. Despite the expectation of some inductive capabilities of ESMFold, we conclude that those are minimal.

## A Supplementary Material

**Figure A1:**
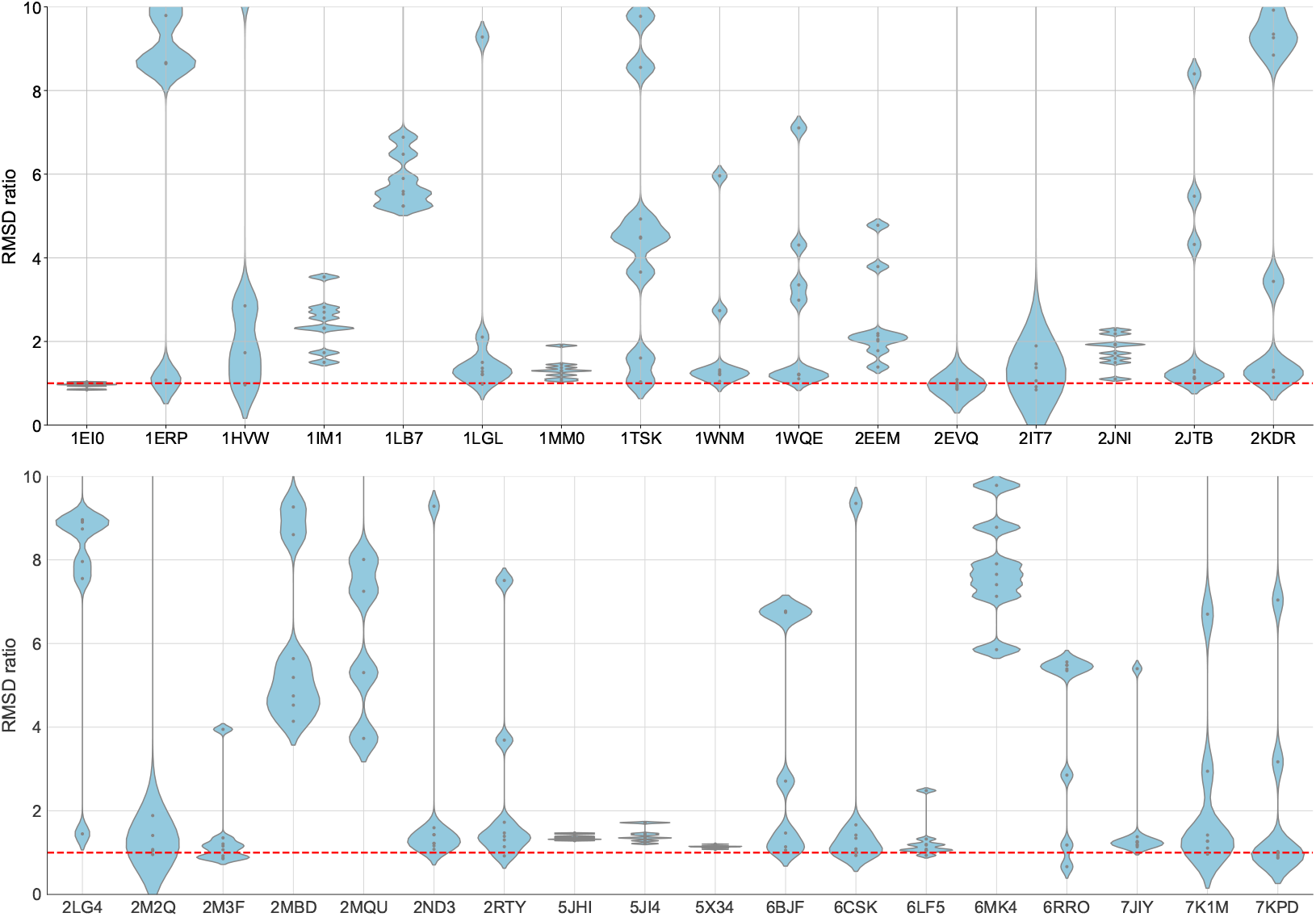
AlphaFold’s prediction accuracy worsens when in the context of a scaffold. Depicted are the ratios of the RMSD of AF2 with respect to the RMSD of AF2 of the tag prediction in isolation. We see that in most cases the ratio is greater than 1 (above the red dashed line) indicating that AlphaFold results in higher error when predicting the tag in context of a scaffold.

**Figure A2:**
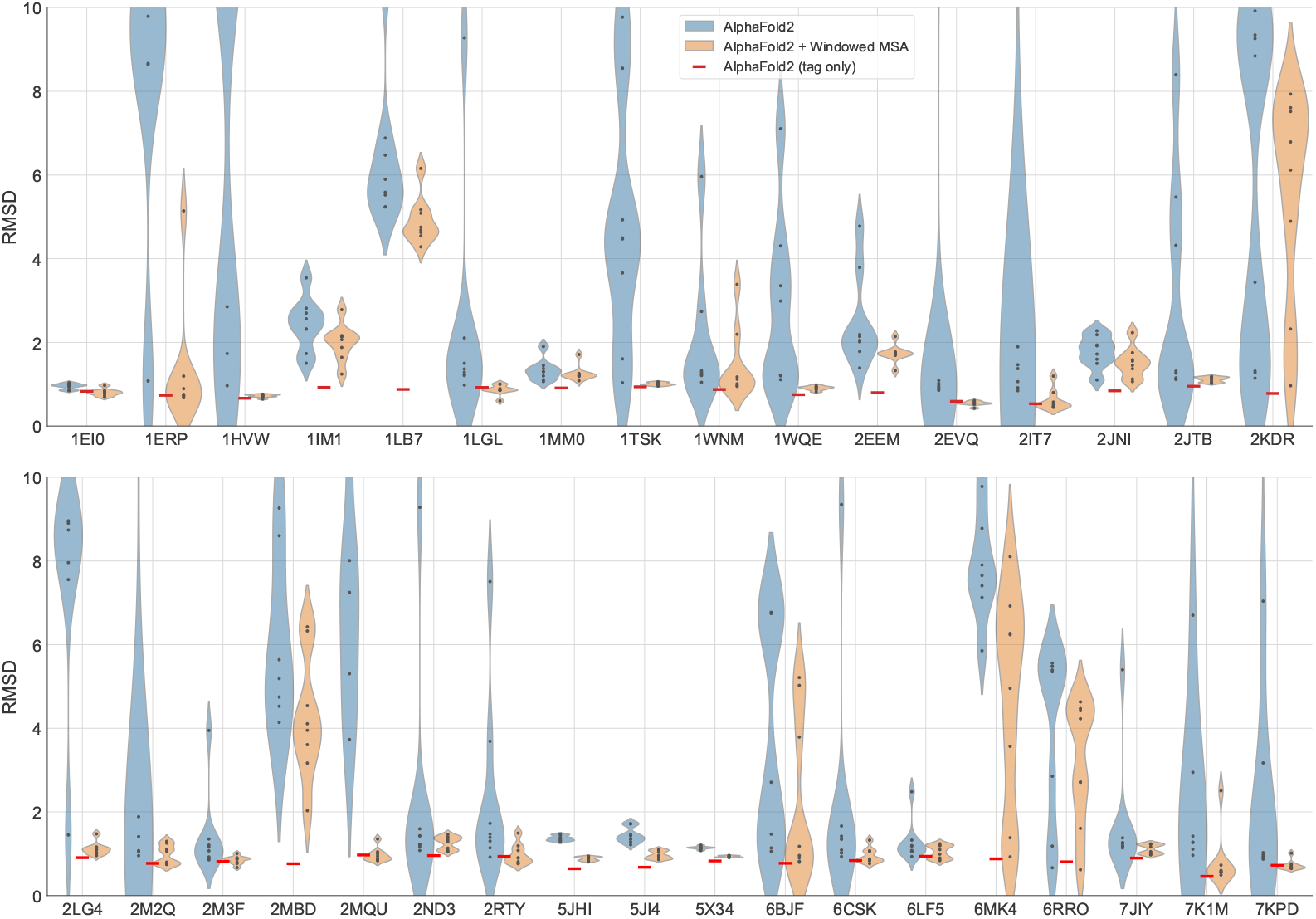
Windowed MSA improves the accuracy of AlphaFold2 predictions. Depicted is the distribution of RMSD (in Å) of each tag predicted in eight different contexts (`N` and `C` chimeras with each of the four scaffold proteins) using AlphaFold2 with and without the proposed windowed MSA “fix”. The accuracy of each tag’s prediction without the scaffold context is shown in red.

**Figure A3:**
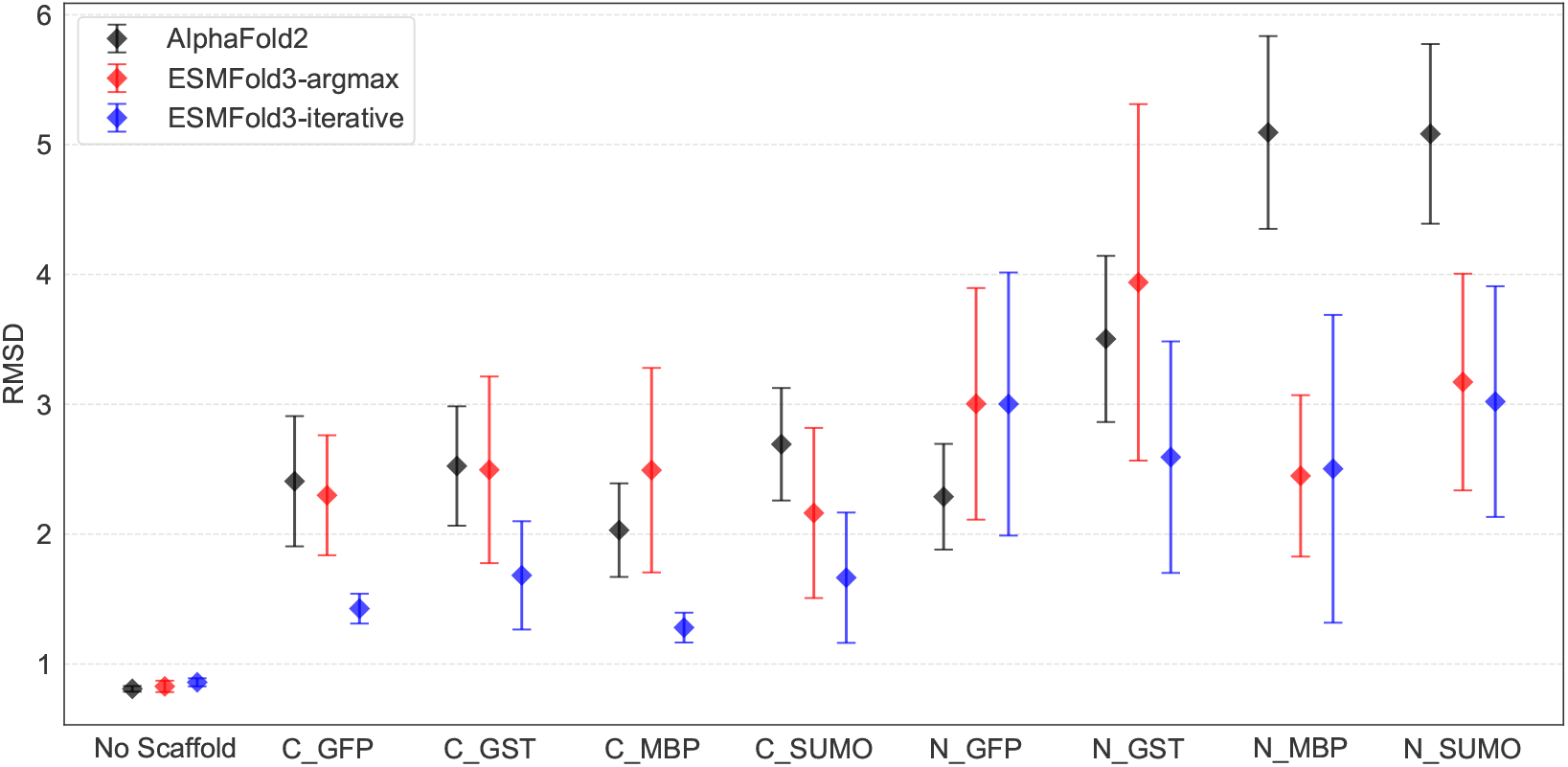
Scaffold context worsens the tag structure prediction accuracy. Note that tags are added as extensions to the native scaffold sequence in all but two cases, `N-MBP` and `N-SUMO` where the tag is replacing part of the native `N` terminal sequence. In these two cases the negative effect of the context is most pronounced

**Figure A4:**
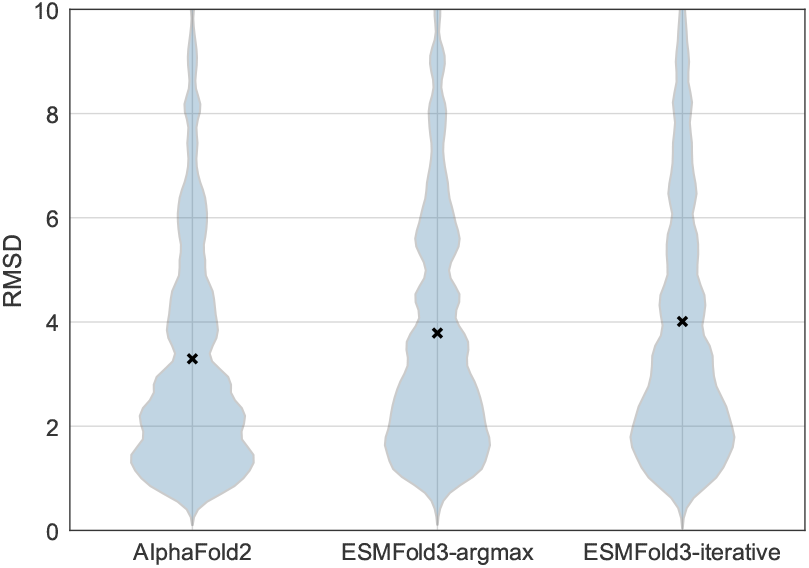
Comparison of the accuracy of AlphaFold2, ESMFold3-argmax, ESMFold3-iterative predictions on the peptide structure prediction benchmark [9].

**Table A1:**
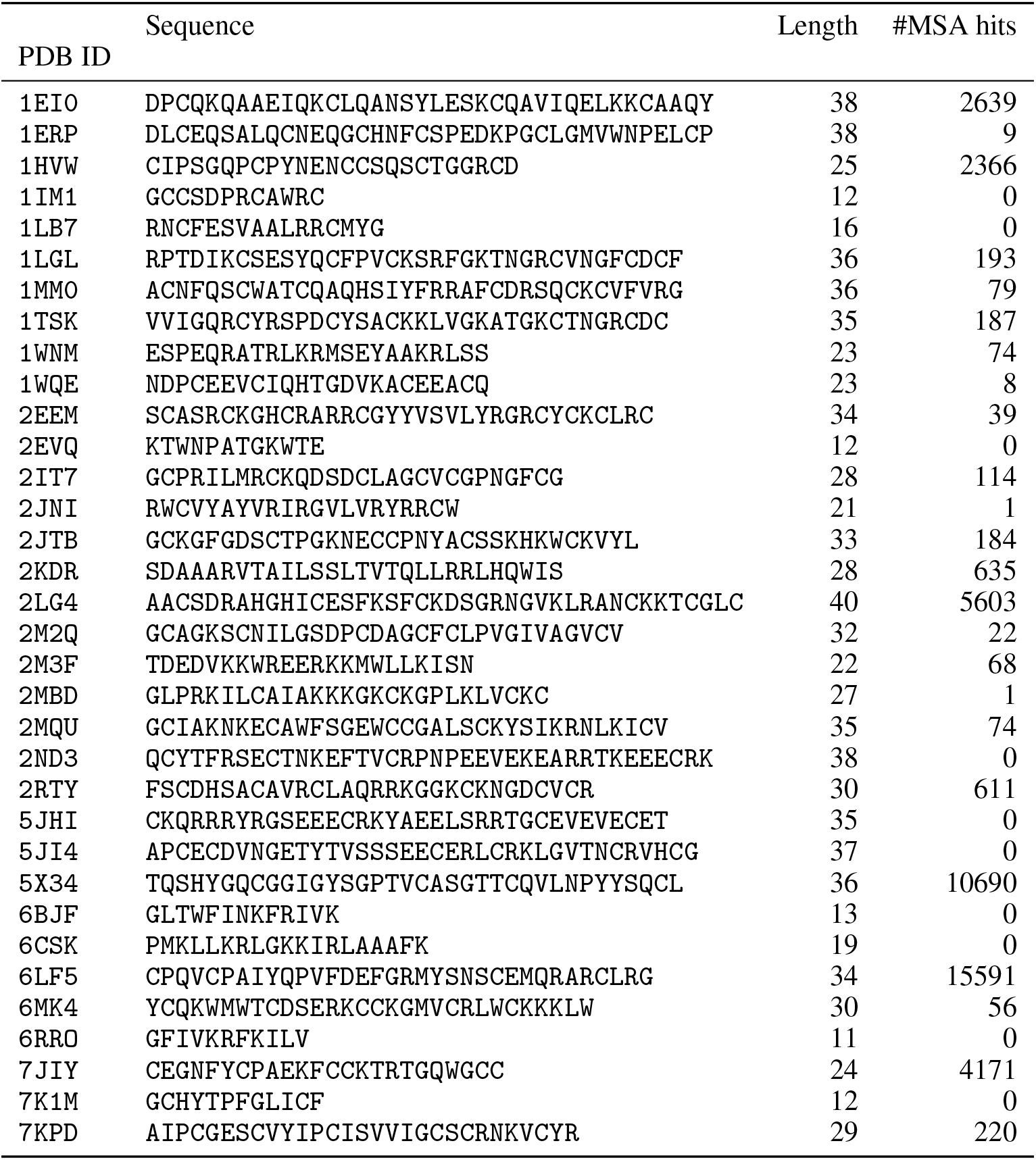
The list of peptide sequences (“tags”) that are considered within this paper.

## Footnotes

1 Accessed on 19 September 2024.

3 This is not entirely true – AlexNet, being a convolutional neural network (CNN), incorporated an important hand-crafted translation-equivariance inductive bias. However, modern visual transformer (ViT) models both outperform CNNs on image recognition tasks and can be claimed to have less hand-crafted biases.

